# Facilitated introgression from domestic goat into Alpine ibex at immune loci

**DOI:** 10.1101/2023.11.23.568345

**Authors:** Xenia Münger, Mathieu Robin, Love Dalén, Christine Grossen

**Affiliations:** Department of Evolutionary Biology and Environmental Studies, University of Zurich, Switzerland; School of Biological Sciences, Monash University, Clayton, Victoria, Australia; Centre for Palaeogenetics, Stockholm, Sweden; Department of Bioinformatics and Genetics, Swedish Museum of Natural History, Stockholm, Sweden; Department of Zoology, Stockholm University, Stockholm, Sweden; WSL Swiss Federal Research Institute, Birmensdorf, Switzerland

**Keywords:** anthropogenic hybridisation, introgression, immune response, ancient genomics, Alpine ibex, Domestic goat

## Abstract

Introgression is expected to be adaptive at loci under balancing selection, for instance at loci relevant for the immune response, but these loci generally also contain ancient polymorphisms. Ancient genomics is a promising tool to detect recent introgression and disentangle it from ancient polymorphisms. Alpine ibex (*Capra ibex*), a wild goat species native to the European Alps, faced near-extinction two centuries ago, but has recovered thanks to successful restoration programs. Previously, signals of introgression from the domestic goat (*Capra aegagrus hircus*) into the Alpine ibex genome were found at the major histocompatibility complex. This introgression was suggested to potentially be adaptive since it introduced new diversity at a gene locus important for immunity. But no genome-wide analyses of introgression have been performed to confirm the potential relevance of immune loci.

Here we take advantage of two ancient whole genomes of Alpine ibex combined with 29 modern Alpine ibex genomes and 31 genomes representing six related Capra species to investigate genome-wide patterns of introgression. Our complementary analysis of putatively introgressed haplotypes and demographic modelling suggests 2.3% recent domestic goat ancestry among wild Alpine ibex. The introgression was estimated to have happened during the last 300 years coinciding with the time when the species had very small population size. Further analysis suggested an enrichment of immune-related genes, where the adaptive value of alternative alleles may give individuals with otherwise depleted genetic diversity a selective advantage.

## Introduction

The transmission of adaptive genetic variation is referred to as adaptive introgression (Hedrick, 2013). The altitude adaptation in Tibetans was reported to be the result of introgression from archaic humans into modern humans (Huerta-Sánchez *et al*., 2014). Another famous example of adaptive introgression was described in mice, where resistance to rodent poison was transferred from the Algerian mouse (*Mus spretus*) to the European house mouse (*Mus musculus domesticus*) (Song *et al*., 2011). Interestingly, introgression was repeatedly shown in genes related to the immune system. Due to pathogen-mediated balancing selection loci of the immune system are usually highly polymorphic (Radwan *et al*., 2020). Hence, receiving variation from a related species may be favorable (Wegner and Eizaguirre, 2012). For example, as modern humans migrated out of Africa, they encountered new environments and locally adapted archaic humans. Accordingly, it was suggested that the bottlenecked migrating human population restored genetic diversity at immune loci after admixture with archaic populations allowing them to adapt to the newly encountered pathogens (Abi-Rached *et al*., 2011; Quach *et al*., 2016; Enard and Petrov, 2018; Gouy, Excoffier and Nielsen, 2020). Exposure to the same environment and hence pathogens may also be involved in the adaptive effect of gene flow at immune-related genes between wild and domestic species (Barbato *et al*., 2017; Chen *et al*., 2018, Howard-McCombe *et al*. 2023). However, the fact that immune-relevant loci are expected to be under balancing selection, also means that some of the variation at those loci is expected to be ancestral polymorphisms (Fijarczyk and Babik, 2015). Ancient genomic data is a promising tool to disentangle ancient polymorphisms (or ancient introgression) from recent introgression (Díez-del-Molino *et al*., 2018).

Alpine ibex, a wild goat species living in high altitude terrain of the European Alps, have recovered after near-extinction, which happened 200 years ago, and are known to hybridize with the closely related domestic goat (Couturier, 1962; Giacometti and Ratti, 2003). During the captive breeding enabling their recovery, Alpine ibex were crossed with domestic goats (*Capra aegagrus hircus)* (Giacometti and Ratti, 2003). Hybrids between the two species are viable and fertile, but their reintroduction was unsuccessful (Couturier, 1962; Giacometti and Ratti, 2003). The failure to reproduce was attributed to the fact that domestic goats and hybrids give birth earlier in the year than Alpine ibex (Bächler, 1918; Couturier, 1962). Nevertheless, hybridization between these species has also been observed in the wild. Feral goats and (suspected) hybrids discovered in the wild are sometimes culled due to the fear of negative effects on the wild population (Moroni *et al*., 2022). However, most Alpine countries do not have clear management regulations in place for dealing with feral goats and hybrids (Moroni *et al*., 2022). A recent survey of 70 populations across the Alps discovered locally very high numbers of suspected hybrids (Moroni *et al*., 2022), many of which have in the meantime been genetically confirmed to be hybrids (personal communication Noel Zehnder).

A previous study reported introgression from domestic goat into Alpine ibex at the major histocompatibility complex (MHC), a gene region important for immune function (Grossen *et al*., 2014). It was suggested that this goat allele may have adaptive effects, since it was found at relatively high frequency in certain populations (Grossen *et al*., 2014). The observation of locally high levels of ongoing hybridisation from a domestic species, combined with potential signals of adaptive introgression, calls for a more in-depth and genome-wide analyses of introgression in Alpine ibex. If the previously reported introgression at the MHC was indeed adaptive, other immune-relevant genes would be expected to show signals of introgression from domestic goat to Alpine ibex. But also potential negative consequences of introgression from a domesticated species are expected and therefore important to be examined. Neither the genome-wide extent of introgression nor the timing(s) of past hybridization event(s) are currently known. It has been hypothesized that the previously described introgression event at the DRB gene happened during the near-extinction of the Alpine ibex (Grossen *et al*., 2014). However, given the shared habitat over centuries, hybridisation and introgression may have already been frequent earlier on.

A major challenge for the study of genome-wide patterns of introgression is to disentangle signals of introgression from signals of ancient polymorphisms, in particular at loci known to be under balancing selection (Fijarczyk and Babik, 2015). As is the case of many other species facing hybridization during or right after a near-extinction event, there are no known non-introgressed Alpine ibex populations, a requirement for many tools used to detect introgression (Sankararaman, 2020). A further challenge to the analysis of introgression among hybridizing species is to disentangle ancient from recent gene flow. Ancient gene flow between ibex species has previously been reported (Pidancier *et al*., 2006).

To alleviate these difficulties, we here combined two ancient whole genome sequences with 29 available modern whole genome sequences of Alpine ibex and 31 whole genomes representing related *Capra* species to analyze genome-wide patterns of recent introgression. We aimed to examine the amount and distribution of genome-wide introgression from domestic goat into Alpine ibex and identify regions with an over– or underrepresentation of introgressed sequences.

## Materials and Methods

### Sample information and sequencing

65 modern, previously published whole-genome sequences representing various goat and sheep species (Grossen *et al*., 2020; NextGen Consortium, https://nextgen.epfl.ch) were analyzed together with two whole-genomes of ancient Alpine ibex individuals. Modern Alpine ibex (*Capra ibex*) were represented by 29 samples from eight Swiss populations and the source of all current Alpine ibex, the Gran Paradiso population in Italy (Figure 1). The other Capra species were represented by four Iberian (*Capra pyrenaica*), two Nubian (*Capra nubiana*) and two Siberian ibex (*Capra sibirica*), one Markhor (*Capra falconeri*), six Bezoar (*Capra aegagrus*) and 16 domestic goat individuals (*Capra aegagrus hircus*) (Figure 1A). Different breeds of domestic goat individuals were included to maximize the genetic diversity and included European, African and Middle Eastern breeds. Additionally, five sheep samples (two *Ovis aries*, one *O. orientalis*, one *O. vignei*, one *O. canadensis*) were included as the outgroup (from here onwards referred to as sheep). One of the two ancient Alpine ibex samples was over 300 years old (1695 CE, based on historical record) and originated from the southern part of Switzerland (Figure 1), the second ancient Alpine ibex sample was over 6500 years old (6590±30BP, based on C14, ETH-62929) and was discovered in a cave in central Switzerland (Robin *et al*., 2022). The genomic data of the domestic goat, Bezoar and sheep are available through the NextSeq Consortium (https://nextgen.epfl.ch). The sequence data of the different ibex species were obtained by Grossen et al. [2020] and Robin et al. [2022] (shotgun sequencing of ancient samples). The ancient samples were USER-enzyme treated before deep sequencing (post-mortem damage was confirmed at the screening step). Therefore, no additional post-mortem assessment was performed for the ancient samples (see Robin et al. [2022] for further detail). The reads were trimmed using Trimmomatic v.0.36 (Bolger *et al*., 2014) and subsequently mapped with bwa-mem v0.7.17 (Li, 2013) to the domestic goat reference genome (ARS1; Bickhart *et al*., 2017). Duplicated reads were marked with MarkDuplicates from Picard v1.130 (http://broadinstitute.github.io/picard). HaplotypeCaller and Genotype GVCF v4.0.8.0 (GATK) were used for the genotype calling and low-quality SNPs were removed with VariantFiltration of GATK. The filter criteria for VariantFiltration were as follows: QD <2.0, FS > 40.0, SOR > 5.0, MQ < 20.0, –3.0 > MQRankSum > 3.0, –3.0 > ReadPosRankSum > 3.0 and AN < 46 (corresponding to 80% of all Alpine ibex individuals in the dataset). We only considered autosomal SNPs (chromosomes 1-29), which had a minor allele count of 1. The obtained autosomal data were further filtered with VCFtools v0.1.16 (Danecek *et al*., 2011). The settings were as follows: Max-missing was set to 0.9, which only retains SNPs with a maximum of 10% missing data and thinning to a minimal among-SNP distance of 100 bp was implemented using the option –-thin 100. Lastly, sites with more than two alleles were discarded (max-alleles 2). This left a final data set consisting of 19,880,663 SNPs across all species.

**Figure 1:**
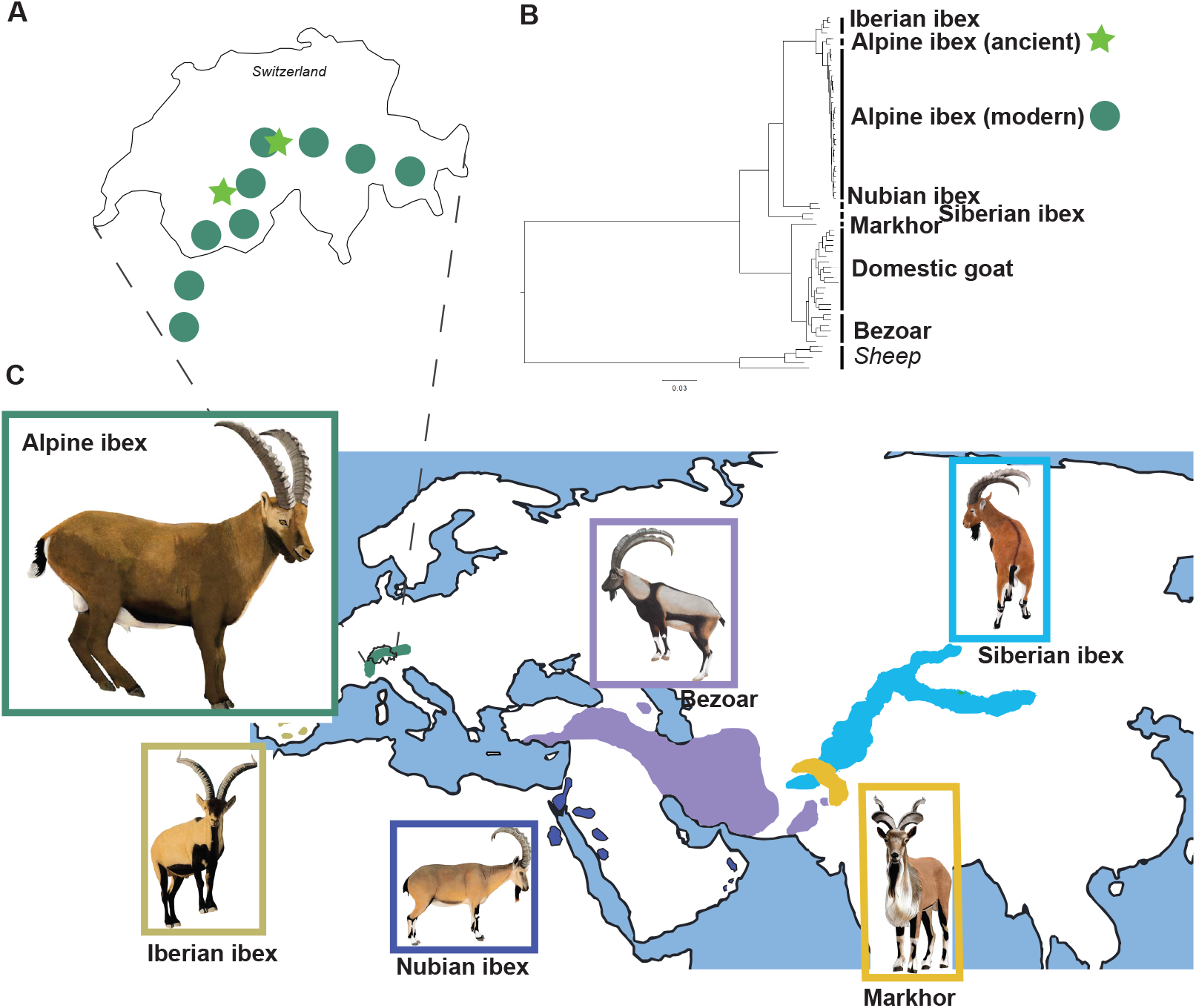
Sample overview. A) Map of Switzerland showing sampling of modern (circles) and ancient (stars) Alpine ibex. B) Phylogenetic tree calculated in RAxML based on autosomal whole-genome data for six *Capra* species and sheep as outgroup. C) Species distribution of five wild *Capra* species used for comparison with Alpine ibex (green).

### Population structure

For a first exploration of the species relationships, we constructed a phylogenetic tree based on maximum likelihood using RAxML v8.2.12 (Stamatakis, 2014). To obtain the input format we used *vcf2phylip* v2.0 (Ortiz, 2019). RAxML was run in the GTRCAT mode. The obtained phylogeny was visualized using FigTree v1.4.4 (Rambaut, 2019).

We further used *ADMIXTURE* v1.3.0 (Alexander *et al*., 2009) to investigate the possibility of recent hybrids among domestic goats and Alpine ibex. VCFtools v0.1.16 (Danecek *et al*., 2011) was used to convert the VCF format into binary PLINK format (bed) for ADMIXTURE. We calculated the cross-validation error for K = {1 to 5}. This indicated that K = 2 is the best number of clusters. Based on this, ADMIXTURE was run for K = {2, 3, 4}. For each K, 15 replicates were calculated.

Principal component analysis was carried out using the R packages {vcfR} v1.10 (Knaus and Grünwald, 2017) and {adegenet} v2.1.2 (Jombart and Ahmed, 2011).

### Genome-wide analyses of admixture

The R package {admixr} v0.7.1 (Petr, 2020), which is an interface for running ADMIXTOOLS v7.0 (Patterson *et al*., 2012) analyses, was used to calculate different F statistics (F_3_, F_4_ and F_4_ ratio). The F_3_ statistic is used either for estimating shared genetic drift between two populations compared to an outgroup or to test whether a population is a mixture between two other populations (Patterson *et al*., 2012). Here, we applied F_3_ statistics for the second purpose. More precisely, we tested whether Alpine ibex are identified as a mixture between domestic goat and another species. A statistically significant negative F_3_ statistic is evidence of admixture. However, a non-significant negative value does not exclude admixture as a possibility. The F_4_ statistic identifies gene flow between populations by examining whether the SNP allele frequency is consistent with an unrooted phylogeny (Patterson *et al*., 2012). We calculated the F_4_ statistic for modern Alpine ibex (A), against ancient specimen or Siberian ibex (B), Bezoar or domestic goat (C) and domestic sheep as the outgroup (D, see also Table 1). A statistically significant deviation from zero indicates gene flow. In a next step, to estimate the goat ancestry among modern Alpine ibex, the F_4_-ratio statistic was calculated by varying the sister taxa of the modern Alpine ibex. Accordingly, the F_4_-ratio statistic took the following form: α = (F_4_(Bezoar, Sheep; modern Ibex, sister taxa))/(F_4_(Bezoar, Sheep; domestic goat, sister taxa)), where the sister taxa was one of the following groups/species: the ancient specimens, the Iberian ibex, Nubian ibex or Siberian ibex (see also Figure 1 and Supplementary Figure S4).

**Table 1:** F_4_ statistics based on different species comparisons (A, B, C, D) and its effect on the detected gene flow (measured as F_4_). Stderr: Standard error of F4 statistics calculated using block jackknife, Z-score: the number of standard errors F_4_ is from 0.

| A | B | C | D | $F_4$ | stderr | Zscore |
| --- | --- | --- | --- | --- | --- | --- |
| **Alpine ibex** | Ancient Alpine ibex | Bezoar | Sheep | -0.00051 | 0.00002 | -25.028 |
| **Alpine ibex** | Ancient Alpine ibex | Domestic goat | Sheep | -0.1145 | 0.00401 | -28.551 |
| **Alpine ibex** | Siberian ibex | Bezoar | Sheep | 0.005245 | 0.000128 | 40.992 |
| **Alpine ibex** | Siberian ibex | Domestic goat | Sheep | 0.2759 | 0.004286 | 64.37 |

### Demographic modelling for inference of genome-wide gene flow

To further investigate the level of potential gene flow between species, we performed demographic modelling using *fastsimcoal2* v2.7.0.9 (Excoffier *et al*., 2013, 2021). *fastsimcoal2* uses site frequency spectra (SFS) to infer demographic parameters via a continuous-time coalescent model. Genotype likelihoods at variable sites were estimated for all autosomes using the samtools model (-GL 1) in ANGSD (Korneliussen, Albrechtsen and Nielsen, 2014). The full site frequency spectra for the derived alleles were calculated using –dosaf with the goat reference genome (ARS1; Bickhart *et al*., 2017) as reference and the ancestral state was defined using sheep as outgroup (-ref and –anc, respectively). A sheep consensus fasta was assembled for this purpose based on a sheep alignment (sample oa.IROA-B2-5037, Table S1) produced using the GAIA pipeline as implemented in Atlas pipeline (Marchi *et al*., 2022) and subsequently performing realignment using GATK v3.7. Next, the tool ANGSD v0.935-53-gf475f10 (option –doFasta 3) was used to create a fasta file.

Reads were filtered for base quality over 20 (-minQ 20) and mapping quality higher than 30 (-minq 30). Furthermore, reads with multiple hits (-uniqueOnly) and bad reads (-remove_bads 1) were removed. ANGSD’s realSFS was used to convert the saf.idx file into the SFS, which was reformatted into *fastsimcoal2* input format using a custom-made script. For each estimated parameter, a predefined range was given, where the lower range limit is an absolute minimum, but the upper range limit is only used to randomly choose an initial value, hence there is no upper limit. The model was set up to estimate long-term effective population sizes for Bezoar (predefined range 2,000 to 2,000,000), domestic goat (predefined range 2,000 to 2,000,000), ghost domestic goat (representing non-sampled goat variation, predefined range 200 to 200,000), modern and ancient Alpine ibex (predefined range 40 to 20,000 and 200 to 2,000,000, respectively). Note that a ghost goat population was introduced because the goat source population is unknown. The model also estimated the time of divergence (informed by previous studies) between the Bezoar and ancient Alpine ibex (predefined range 80,000 to 250,000 generations; Manceau *et al*., 1999), Bezoar and domestic goat (predefined range 626 to 45,000 generations; domestication latest 5000 years ago assuming a generation time of 8 years, Luikart *et al*., 2001), ancient and modern Alpine ibex (predefined range 112 to 1,600 generations; Pidancier *et al*., 2006; Ureña *et al*., 2018) and domestic goat and ghost domestic goat (predefined range 101 to 625 generations). Additionally, the model also estimated the rate and timing of gene flow between the ghost domestic goat and modern Alpine ibex. The predefined range of the long-term gene flow was set to 0 to 25% in both directions. Furthermore, the start of recent gene flow (predefined range 9 to 100 generations; Grossen *et al*., 2014) and end of recent gene flow (predefined range 0 to 8 generations; Grossen *et al*., 2014) were added to the model. In *fastsimcoal2*, to estimate the frequency spectra through simulations, the data type was set to FREQ. With these input settings, we performed 100 independent simulation runs each computing the SFS for the derived alleles in the population (-d) with 200 000 simulations for each set of parameters (-n 200 000) with 50 loops of parameter estimation (-L 50) by maximum composite likelihood (-M). All runs were ordered according to smallest AIC and highest maximum likelihood value. According to the ranking we took the mean for each parameter of the 10 best runs. We followed the *fastsimcoal2* manual to generate non-parametric bootstrapped SFS. This step included modification of the parameter input file (.par) to model 200 000 non-recombining independent loci with the data type DNA of a length of 1000 bp. The modified. par file was subsequently used in *fastsimcoal2* to create 100 SFS. This step included the options –n1 (one simulation performed for each parameter file), –b99 (number of bootstraps performed on polymorphic sites), –s0 (output all SNPs along a DNA sequence) and –l (generating mutations according to an infinite site mutation model). As a last step, *fastsimcoal2* estimated the parameters based on these bootstrapped SFS with the settings: 30 loops performed to estimate the parameters (-L) by maximum composite likelihood (-M), 100 000 simulations per parameter file (-n), computing the SFS of the derived alleles for each population sample (-d). Based on these bootstrap runs the 95 percentile confidence intervals were calculated.

### Admixture analyses along the genome

We expected varying levels of introgression along the Alpine ibex genomes. To infer the most likely origin (either museum specimen or domestic goat) of a certain genomic position in an Alpine ibex genome we used the tool admixfrog 0.6.2.dev6 (Peter, 2020). Admixfrog compares allele frequencies at each genomic position in a target (here Alpine ibex) to a number of sources (here either ancient Alpine ibex or domestic goat). A bin size of 10,000 bp was chosen and we used a physical map. The obtained information was further analyzed using custom scripts in R v3.5.3 (R Core Team, 2018).

### GO term identification of introgressed segments

To identify possible gene enrichments in introgressed regions, an analysis of gene ontology (GO) terms was performed. We searched the annotated coding sequences within regions identified as introgressed by admixfrog for overrepresentation of biological processes, molecular functions and cellular components. A significance level p = 0.05 was set. Since previous introgression was found at the immune system, specifically at the MHC, we also carried out a GO term analysis excluding coding sequences annotated as MHC and another analysis excluding MHC, immune and antigen. This allowed us to identify other potentially enriched gene categories among introgressed regions. For the GO term analysis the following R packages were used: {annotate} v1.56.2 (Gentleman, 2018), {GO.db} v3.5.0 (Carlson, 2017), {GSEABase} v1.40.1 (Morgan, Falcon and Gentleman, 2017), {GOstats} v2.44.0 (Falcon and Gentleman, 2007).

### R packages

For multiple format conversions as well as visualization we used the software R v3.5.3 (R Core Team, 2018) with the following packages: {admixr} v.0.7.1 (Petr, 2020), {annotate} v1.56.2 (Gentleman, 2018), {ggplot2} v3.3.2 (Wickham, 2016), {ggpubr} v0.4.0 (Kassambara, 2020), {ggrepel} v0.8.2 (Slowikwski, 2020), {GO.db} v3.5.0 (Carlson, 2017), {GOstats} v2.44.0 (Falcon and Gentleman, 2007), {GSEABase} v1.40.1 (Morgan, Falcon and Gentleman, 2017), {splitstackshape} v1.4.8 (Mahto, 2019), {svglite} v2.0.0 (Wickham *et al*., 2020) and {tidyverse} v1.3.0 (Wickham *et al*., 2019).

## Results

To investigate genome-wide patterns of introgression from domestic goat to Alpine ibex, a total of 67 samples consisting of eight different species were used. This included whole-genome sequence data from 29 modern Alpine ibex covering a large part of the current home range by representing eight Swiss populations and the source of all current Alpine ibex populations, the Gran Paradiso population in Italy, two ancient specimen of Alpine ibex (both from Switzerland, about 300 and 6,500 years old) and 16 domestic goats (Figure 1). Additionally, 15 specimen representing related *Capra* species (including 6 Bezoar, the direct ancestor of domestic goat) were analysed (Figure 1C). The final data set included 19,880,663 SNPs. For the Bezoar and domestic goat the mean read coverage was 12X (6X-19X), the two ancient specimen of the Alpine ibex had a mean coverage of 11X (8X and 14X) and for the other species the mean coverage was 18X (8X-38X).

### Clear species distinction between Alpine ibex and domestic goat

For an initial overview of the genome-averaged species relationships including the relation of ancient and modern Alpine ibex, a phylogenetic tree was constructed using RAxML (Stamatakis, 2014). As expected from previous analyses and supported by a PCA across species (Supplementary Figure S1) (Pidancier *et al*., 2006; Grossen *et al*., 2020; Robin *et al*., 2022); individuals clustered by species. The two ancient Alpine ibex specimens built a separate branch basal to modern Alpine ibex (Figure 1B). This confirms previous results based on mitogenome data suggesting that these two specimens represent lineages which disappeared during the near-extinction (Robin et al. 2022).

Next, we wanted to exclude the possibility that the modern Alpine ibex contained any early hybrids, because this would confound the introgression analysis. Both analyses, the Principal Component Analysis and the clustering in ADMIXTURE (Alexander *et al*., 2009) suggested that the data set did not include any early generation hybrids (i.e. F1 or backcross, Figure 2). In the PCA the first principal component discriminated the domestic goats from the Alpine ibex, with no intermediate specimen (which would be expected for early hybrids) as well as the modern from the ancient samples of the Alpine ibex (Figure 2A). The second principal component distinguished the geographic origins of the domestic goats (especially the European breeds from Moroccan and Iranian, Figure 2A). The clear discrimination between domestic goat and modern Alpine ibex was also reflected in the ADMIXTURE results (most likely number of groups K = 2; Figure 2B, Supplementary Figures S2 and S3). The assignment of the domestic goat and Alpine ibex samples to the two groups always reflected the corresponding species. The ancient Alpine ibex were assigned to the same group as the modern Alpine ibex. Only a small proportion of the ancient Alpine ibex was assigned to the same group as domestic goat (Figure 2B) likely reflecting the higher diversity among the ancient Alpine ibex than modern Alpine ibex recently reported based on mitogenome analyses (Robin *et al*., 2022). Increasing the number of groups (K) identified population structure among domestic goats for K=3 (separating European from Iranian and Moroccan goats) and at K = 4, the ancient Alpine ibex specimens were assigned to a distinct group in 12 out of 15 replicates (Supplementary Figure S2).

**Figure 2:**
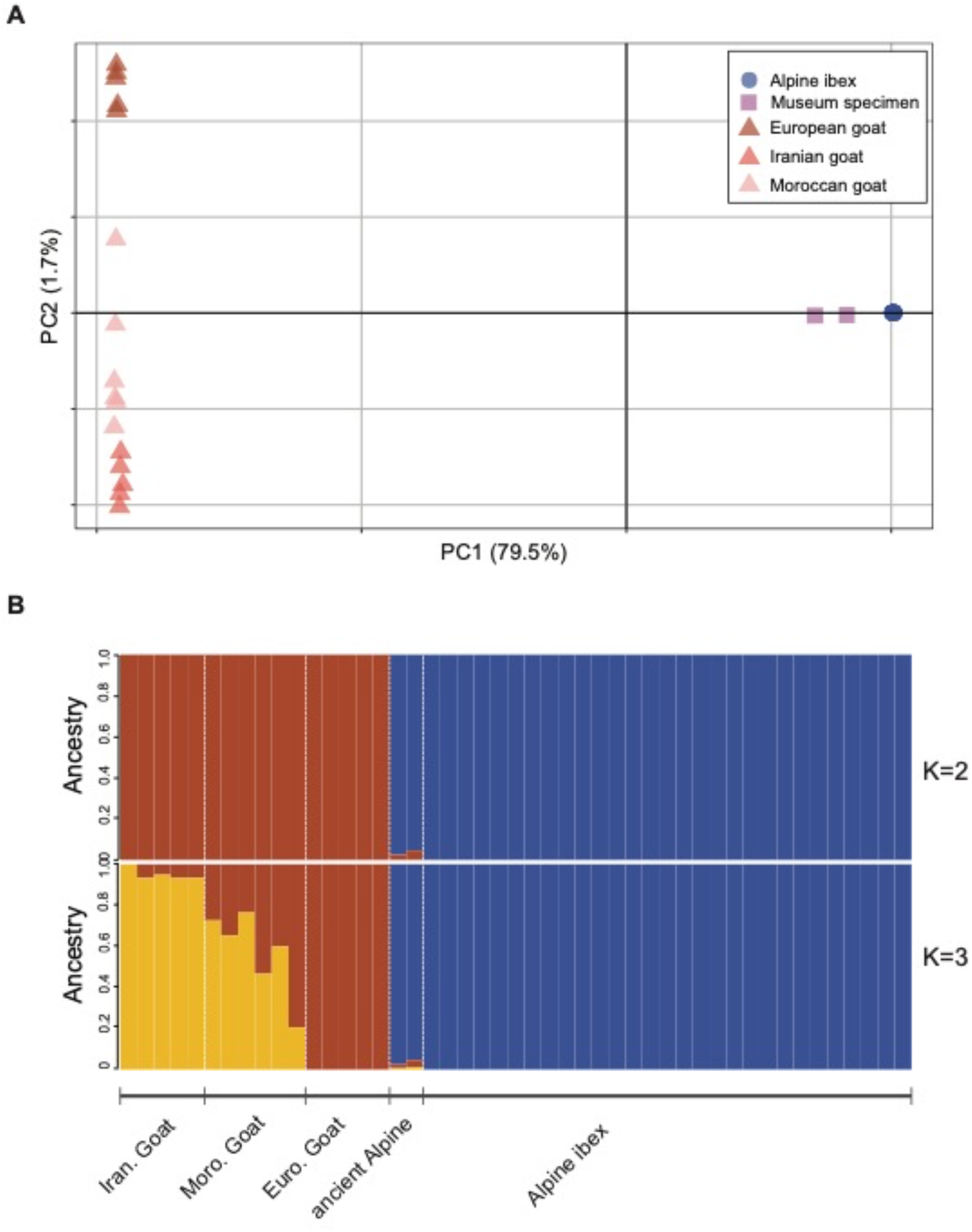
Alpine ibex and Domestic goat. (A) Principal Component Analysis of the modern (circles) and ancient (rectangles) samples of Alpine ibex and domestic goat samples (triangles) representing European, North African and West Asian breeds. (B) ADMIXTURE plots for K=2 and K=3. The domestic goats are distinct from modern Alpine ibex. Best cross validation error for K=2 (see also Supplementary Figures S2 and S3).

### Low rates of admixture among domestic goat and modern Alpine ibex

PCA nor ADMIXTURE identified any early-generation hybrids among the sequenced individuals. To explore potential low levels of admixture, we took advantage of F statistics (Reich *et al*., 2009; Patterson *et al*., 2012).

First, F_3_ statistics were used to test if Alpine ibex can be considered as an admixed population of domestic goat and another group (See Supplementary Table S2). None of the tested species trios resulted in a significant negative value that would suggest admixture. Hence, no evidence of admixture between the tested groups was found (Table S2). Importantly, the absence of admixture evidence with the F3 statistics does not allow the conclusion that no admixture between the groups has happened (Peter, 2016). F_4_ statistics were used to test for admixture in an alternative way (Table 1). Also F_4_ statistics compare allele frequencies but instead of only three populations, four populations are used and the forth population is usually a divergent outgroup (here sheep), from which no admixture into any of the other is expected (Patterson *et al*., 2012). When using the two ancient specimens of Alpine ibex basal to all modern Alpine ibex (1695 CE and 6590±30BP) as the sister group to modern Alpine ibex, we obtained significant negative values suggesting gene flow between the ancient specimens and Bezoar/domestic goat, while testing against the Siberian ibex resulted in significantly positive values suggesting gene flow between Bezoar/domestic goat and Alpine ibex (Table 1).

Finally, to estimate admixture proportions α from domestic goat (B) for each Alpine ibex population (X), F4-ratios were calculated, where α = F_4_(A, O; X, C) / F_4_(A, O; B, C) (Patterson *et al*., 2012, see also Supplementary Figure S4A). The calculated admixture proportion varied considerably. The F_4_-ratio estimation with C = ancient specimen yielded a mean admixture proportion of –0.83% domestic goat ancestry across the modern Alpine ibex populations. With C = Iberian ibex the mean admixture proportion was –0.1% domestic goat ancestry. With C = Nubian ibex the mean admixture proportion was 0.26% domestic goat ancestry and with C = Siberian ibex the admixture proportion was 7.7% domestic goat ancestry (Supplementary Figure S4B). Hence, estimates of domestic goat ancestry proportions in Alpine ibex increased with phylogenetic distance from (and decreased shared drift with) the sister group used (Figure 1).

### Estimating admixture proportions using demographic modelling

Our F statistics analyses revealed a strong effect of the chosen sister group and F-statistics possibly due to varying levels of shared drift and (ancient) gene flow among the test populations. To take divergence times and effective population size (both defining factors for shared drift) in account by estimating them together with gene flow rates, we chose a demographic modelling approach using *fastsimcoal2.* Coalescent simulations with *fastsimcoal2* estimated the long-term effective population size of diploid individuals for Bezoar to 42,083 (95% percentile CI: 41,262-43,649), for domestic goat to 4,185 (95% percentile CI: 1,350-4,775), for the ghost goat population (assumed non-sampled source of admixture) to 39,863 (95% percentile CI: 13,320-93,387), for the modern Alpine ibex to 32 (95% percentile CI: 20-185) and for the ancient Alpine ibex to 201,666 individuals (95% percentile CI: 196,207-200,688; Figure 3). The split time between Bezoar and ancient Alpine ibex was estimated to 655.3 kya (95% percentile CI: 647.1-672.9 kya), between Bezoar and domestic goat to 39.3 kya (95% percentile CI: 11-44.0 kya), between domestic goat and ghost goat to 2.5 kya (95% percentile CI: 1.3-5.0 kya) and between ancient and modern Alpine ibex to 6.4 kya (95% percentile CI: 1.6-11.6 kya; Figure 3). Gene flow from ghost goat into modern Alpine ibex was estimated to 2.3% (95% percentile CI: 1.2-21%), while the opposite direction was estimated to 0.8% (95% percentile CI: 0.6-4.7%). Furthermore, the gene flow was estimated to have started 305 years ago (∼42 generations, 95% percentile CI: 128-763.4 years ago, Figure 3) and ended 0.8 years ago (95% percentile CI: 0-11.8 years ago; Figure 3).

**Figure 3:**
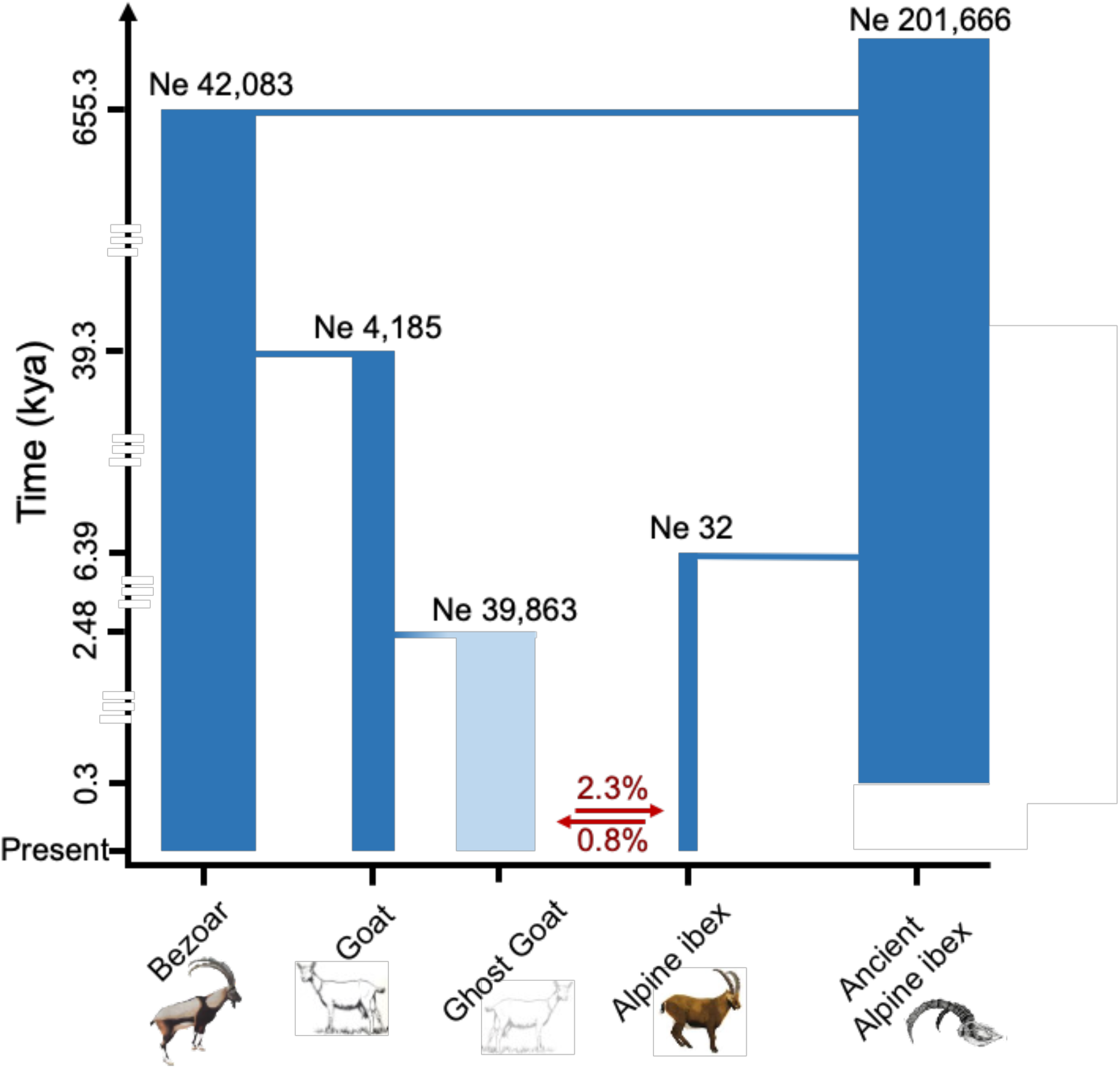
Demographic model estimating split times, long term effective population size and gene flow between ghost goat and Alpine ibex using coalescent simulations in *fastsimcoal2* (Excoffier *et al*., 2021). Light color of ghost goat indicates that there were no samples available for this group and it was simulated as non-sampled variation.

### Long-term species-wide gene flow estimates reflect inferred goat ancestry in modern Alpine ibex

Based on the demographic model described above, the species-wide rate of gene flow from domestic goat (modeled as non-sampled ghost goat population) into Alpine ibex was estimated to 2.3%. Next, we were interested in how much introgressed material each Alpine ibex individual carries. To this end, we implemented the tool admixfrog (Peter, 2020) which compares the allele frequency at each genomic position in a target to a number of sources (here either ancient Alpine ibex or domestic goat). The total sequence length in heterozygous state ranged from around 32 Mb to 52 Mb and between 32 Mb to 42 Mb in the homozygous state (Supplementary Figure S5). Thus, the mean percentage of goat ancestry found among the Alpine ibex individuals was 2.43% with relatively small among-individual variation.

### Gene ontology term analyses of introgressed loci

Multiple generations after the initial hybridization the introgressed sequences are expected to be broken up into smaller pieces by recombination. Additionally, alleles located on the introgressed sequence interact with the other genes. This may positively or negatively affect the individual fitness. Previous studies in various organisms have shown adaptive introgression especially at genes of the immune system (Enard and Petrov, 2018; Fijarczyk *et al*., 2018; Dudek *et al*., 2019). To investigate the potential accumulation of putatively introgressed sequences at genes with a certain biological function, gene ontology (GO) term analysis was carried out. The regions identified as introgressed by admixfrog overlapped with 496 genes across all autosomes. The detected regions were not uniformly distributed across autosomes (Figure 4, Supplementary Figure S6). The highest number of overlaps was found on chromosome 23 (over 70 overlaps, see also Figure 4) and chromosome 15 (over 40 overlaps) while less genes overlapped with putatively introgressed regions on for instance chromosomes 14, 21 and 22. On chromosome 23, a large number of overlaps occurred with the MHC and other immune-related genes. The GO term analysis suggested a strong and highly significant enrichment for introgressed regions in the categories of response to stimulus (p < 0.0001, 319 of 4625 genes), antigen processing and presentation (p < 0.0001, 20 of 48 genes) and immune response (p < 0.0001, 60 of 431 genes, Figure 5). Since these results are expected to be driven by the MHC complex situated on chromosome 23 (Dong et al. 2012), the same analysis was performed by excluding all MHC associated coding sequences or all immune associated coding sequences (including the terms “MHC”, “immun” or “antigen” in the annotation), respectively (Supplementary Figure S7). We still observed several immune-related biological processes enriched after excluding all MHC associated coding sequences (Supplementary Figure S8), but the most significant enrichments were now for G-protein coupled receptor signaling pathway (p < 0.0001, 133 of 1458 genes), response to stimulus (p > 0.0001, 300 of 4311 genes) and cell communication (p > 0.0001, 247 or 3562 genes).

**Figure 4:**
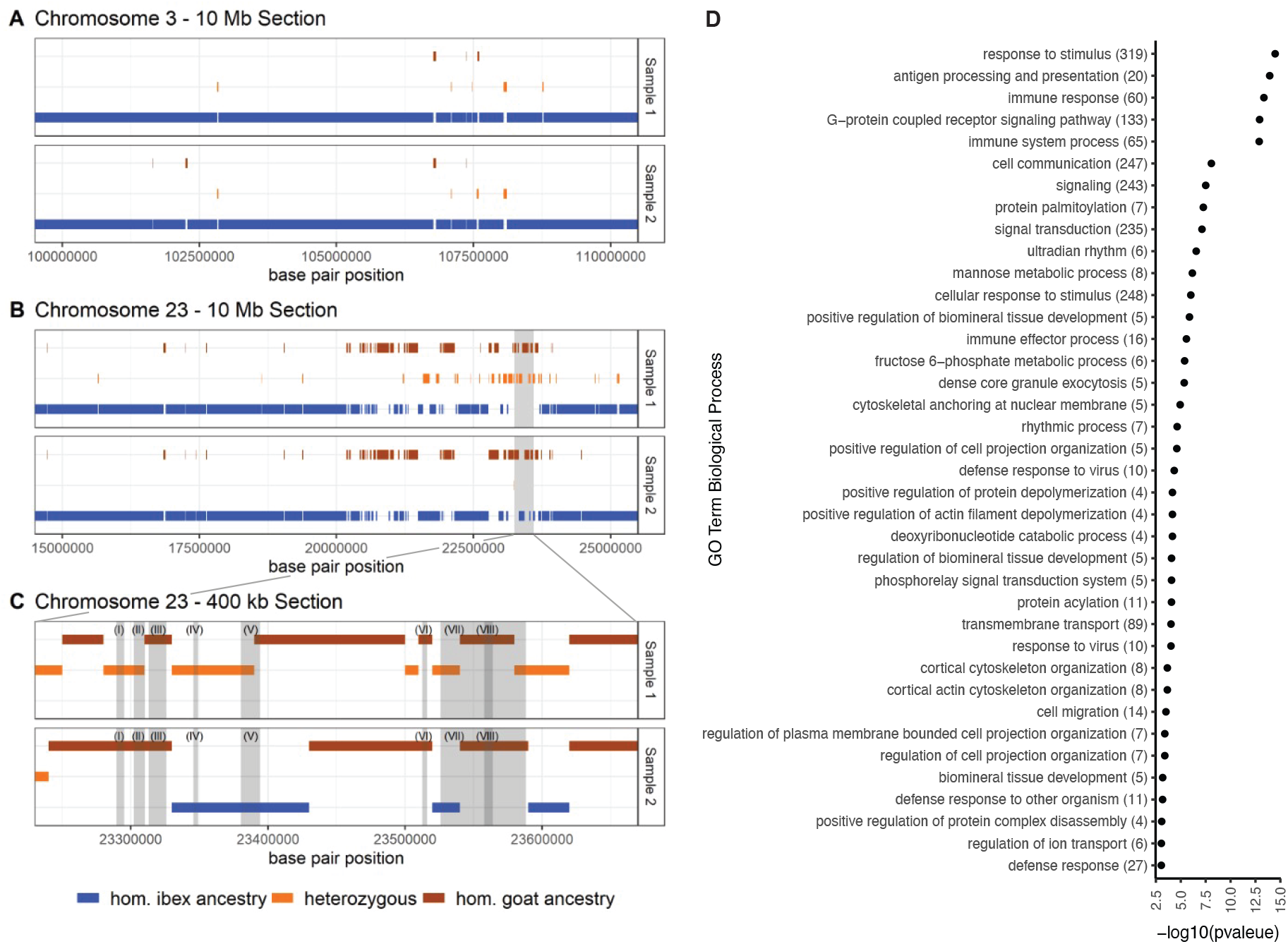
Analysis of introgressed haplotypes. A) A 10 Mb section of chromosome 2 of two Alpine ibex individuals illustrating the admixfrog ancestry inference, where only little introgressed sequences were found. B) A 10 Mb section of chromosome 23 with many introgressed sequences found. Both individuals show stretches of homozygous introgressed material. One individual shows also clear signals for heterozygous introgressed sequences. The grey shading highlights the enlarged part in (C). C) A 400 kb section of chromosome 23, where the grey shading indicates annotated exons. The labelled exons are: (I) butyrophilin-like protein 1, (II) butyrophilin subfamily 1 member A1, (III) butyrophilin-like 2, (IV) HLA class II histocompatibility antigen DR α-chain (transcript variant X1), (V) DLA class II histocompatibility antigen DR-1 β-chain-like, (VI) HLA class II histocompatility antigen DRB1 4-β-chain, (VII) SLA class II histocompatibility antigen DQ haplotype D α-chain, (VIII) boLa class II histocompatibility antigen DQB β-chain (transcript variant X3) and (IX) HA25. D) GO term summary for biological processes of introgressed sequences detected by admixfrog. Numbers in brackets specify the number of genes in each category. Colors indicate log10 p-values of enrichment (lower p-values for darker blue).

Next, we were interested in what GO terms were underrepresented among introgressed haplotypes as an indicator for gene categories putatively under negative selection if introgressed. Among the ten most significant hits, nine were metabolic processes and gene expression (Supplementary Figure S8), all processes expected to be under purifying selection due to their conserved nature (Wilson *et al*., 1977; Kimura, 1983).

## Discussion

High variation at immune system genes has been proposed to improve the survival of individuals (Sommer, 2005) and these genes are generally assumed to be under balancing selection (Croze *et al*., 2016). Theory predicts that new alleles originating from mutations or introgression are favored in genomic regions under balancing selection (Schierup *et al.,* 2001; Castric *et al*., 2008). Targeted sequencing in allopatric and parapatric newts supports the prediction that balancing selection facilitates introgression (Fijarczyk *et al*., 2018). It was therefore suggested that introgression may be an important process shaping the high variation found at the MHC, a major gene complex of the immune system (Wegner and Eizaguirre, 2012) and immune-relevant loci in general (Reilly *et al*., 2022). Adaptive introgression at the MHC has indeed been suggested in a number of species including humans (Abi-Rached *et al*., 2011), newts (Gaczorek *et al*., 2023) and lizards (Gaczorek *et al*., 2023). The generally high genetic diversity at the MHC loci in domestic livestock was suggested to originate from intentional or accidental backcrosses with their wild ancestors (Vilà et al., 2005). But evidence from the wild remains scarce and is still mostly based on a limited number of markers. Even less studies investigated possible adaptive introgression at immune-relevant genes in general. Evidence for adaptive introgression at immune-relevant genes comes most of all from humans (Quach *et al*., 2016; Enard and Petrov, 2018; Gouy and Excoffier, 2020), a study based on a SNP-chip suggested adaptive introgression at immune-relevant gene regions from mouflon to domestic sheep (Barbato et al. 2017) and a very recent study on wildcats (Howard-McCombe *et al*. 2023).

The whole genome sequences of the modern Alpine ibex revealed clear differentiation from the domestic goat samples indicating that no early generation hybrids (e.g., F1, backcross individuals) were included in this analysis. However, the demographic model suggested low genome-wide levels (2.3%) of introgression from domestic goat into Alpine ibex. Under the assumption of neutral evolution, this level of introgression is in accordance with the main introgression event having taken place between 5 to 6 generations ago (3.1 to 1.6 percent). Assuming a generation time between 8 and 10 years (Stüwe and Grodinsky, 1987), the main hybridization event leading to this level of introgression would have happened only 48 to 60 years ago rather than during the near extinction of Alpine ibex as was proposed previously (Grossen *et al*., 2014). Yet the start of gene flow was estimated to much earlier, about 300 years ago. And the GO term analysis identified an enrichment of immune-relevant genes overlapping with the introgressed regions. The discrepancy between estimated time period and amount of introgression together with the enrichment analysis suggests that the introgressed haplotypes did not evolve neutrally but rather were preferentially retained at immune-relevant genes while potential negative selection acted in other gene regions.Our F-statistics outcomes were highly sensitive to the chosen sister group with an apparent correlation with phylogenetic relationship. If a closely related population or species was used as the sister group (i.e., ancient specimen or Iberian ibex), gene flow was suggested to be more likely between domestic goat and either of the two groups or between Alpine ibex and domestic sheep. However, for the two more distantly related species (Nubian and Siberian Ibex) relatively high proportions of domestic goat ancestry were estimated in Alpine ibex. It is worth noting that Iberian ibex also experienced a strong bottleneck in the recent past and hybrids with domestic goat have been reported repeatedly (Cardoso *et al*., 2021). It is therefore possible that Iberian ibex received gene flow from domestic goat in the past and the Nubian and Siberian ibex received less. However, it is less straight-forward to explain the apparent admixture between ancient Alpine ibex and domestic goat except that potentially the more recent specimen of the two, dating to 1695 already received some introgression. Therefore, these results are to be interpreted with caution. Possible effects of gene flow into ancient Alpine ibex is also reflected in the estimates of effective population size for ancient Alpine ibex, which with over 200’000 were very high and more likely an effect of population structure (due to both space and time difference between the two ancient samples) and potential gene flow from related species rather than very large populations in the past. Hence, the estimate of 2.3% domestic goat ancestry in modern Alpine ibex is an estimate of recent introgression, while overall gene flow between the two species was potentially higher. The small Ne estimate for modern Alpine ibex (Ne=32) on the other hand is a likely result of the near-extinction.

Due to this history of near-extinction, Alpine ibex have a low genome-wide diversity (Grossen *et al*., 2018; 2020). Our genome-wide analysis suggesting that introgressed haplotypes are more likely retained at immune-relevant genes is in accordance with introgression potentially being beneficial at such loci. Higher MHC-diversity was indeed recently shown to correlate with higher resistance to keratoconjunctivits (an infection of the eye) in Alpine ibex (Brambilla *et al*., 2018) and another study found an association between two immune-relevant loci (including Toll-like receptor) and the risk of brucellosis infection (Quéméré *et al*., 2020). However, regions under balancing selection are candidates for ancient polymorphisms and disentangling signatures from introgression and ancient polymorphisms are not trivial. Our approach of using two ancient genomes as baseline alleviates this issue by focusing the analysis on recent gene flow.

We were also interested in genomic regions potentially under negative selection. We found that genes involved in metabolic processes and gene expression were underrepresented among putative introgressed haplotypes. These genes are expected to be under purifying selection. However, although introgression is likely to be disfavored under purifying selection, also the power to detect introgression is lower due to the generally low divergence in such regions (Schumer *et al*., 2016).

In conclusion, introgression may affect a species on a wide spectrum of outcomes (Edmands, 2007). Adaptive benefits of introgression under certain selection schemes may drive the retainment of introgressed sequences as suggested by this study for immune-relevant loci. Potential positive effects of adaptive introgression and genetic rescue can increase adaptive genetic variation and for instance improve adaptation to environmental changes (Brauer *et al*., 2023) or resistance to disease (e.g., Enard and Petrov, 2018; Gouy *et al*., 2020, Howard-McCombe *et al*. 2023 and this study). However, this is context-dependent and introgression may be maladaptive in another selective environment (Dolgova and Lao, 2018; Zeberg and Pääbo, 2020). Hybridisation is a major concern and subject of debate in conservation biology (Allendorf *et al*., 2001; Quilodrán *et al*., 2020). It is observed with particular attention if it happens with domesticated species as for instance in wildcats (Howard-McCombe *et al*., 2021) or wolves (Muñoz-Fuentes *et al*., 2010; Galaverni *et al*., 2017). Anthropogenic introgression can be a serious threat for a species potentially leading to negative fitness effects (e.g. plains bison, Hedrick, 2010) up to swamping as seen in the Scottish wildcat (Senn *et al*., 2019; Howard-McCombe *et al*., 2021). To inform the conservation management of species, a better understanding of hybridization and its consequences in the wild is required. Improved understanding of the balance between positive versus negative effects of hybridisation and how this affects the faith of introgressed gene regions in the wild (i.e., how likely introgressed gene regions will be retained over multiple generations), would be particularly informative. In the case of the Alpine ibex, the introgression is mainly localized at genes of the immune system. Thus, at least in this species, the introgression from the domestic goat into its wild, inbred relative may have had some positive effects. As previously mentioned, the outcome cannot be anticipated, due to the interplay of many factors. Hence, the common practice of quickly removing hybrids resulting from anthropogenic hybridization from the wild gene pool is the most conservative approach, even if this may represent a missed chance to increase the genetic variation of an otherwise genetically depleted species.

## Acknowledgments

This project was funded by the Swiss National Science Foundation (31003A_182343 to C.G.). This study makes use of data generated by the NextGen Consortium (grant number 244356) of the European Union’s Seventh Framework Program (FP7/2010-2014). The authors also acknowledge support from the Science for Life Laboratory, the National Genomics Infrastructure and UPPMAX for providing assistance in massive parallel sequencing and computational infrastructure.

## Data Accessibility and Benefit-Sharing

The two ancient whole genome sequences used for this project and corresponding Metadata will be deposited at the NCBI Short Read Archive under the Accession nos. XX and YY (BioProject XYZ).

## Author Contributions

C.G. conceived the project. M.R. acquired the ancient samples and carried out the sample and library preparation. X.M. conducted all bioinformatic analysis with input from M.R. and C.G.. X.M. and C.G. wrote the manuscript. L.D. provided access to an ancient laboratory, consumables, and sequencing facilities for data generation of the two ancient specimens.

