## Supplementary Material for "Facilitated introgression from domestic goat into Alpine ibex at immune loci"

Münger et al.

### Supplementary Figures

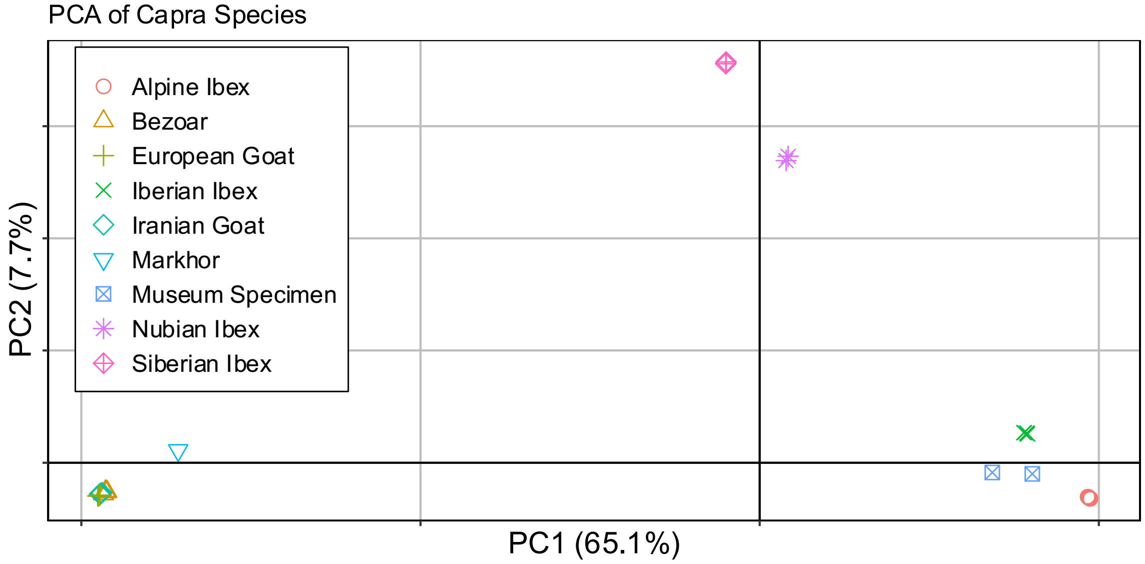


**Supplementary Figure S1**: Principal Component Analysis visualizing genetic distinctiveness of six Capra species.

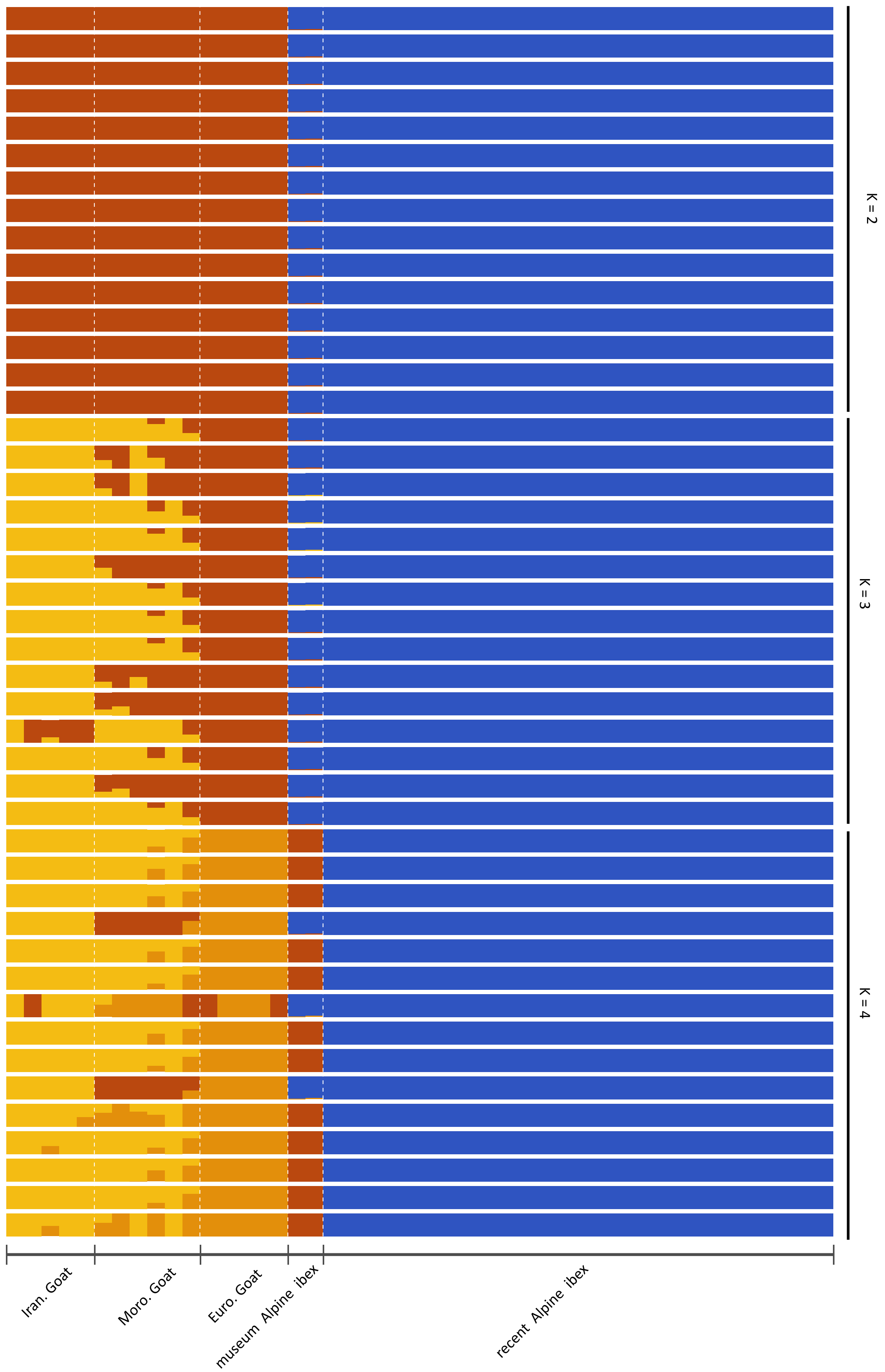


**Supplementary Figure S2**: Admixture results for K = 2, 3 and 4 with 15 replicates each. Best cross validation error for K=2.

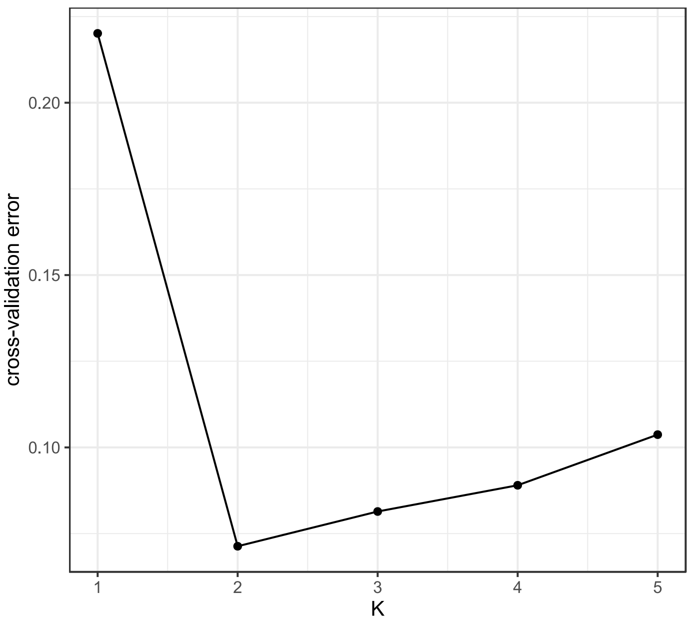


**Supplementary Figure S3:** Cross-validation of ADMIXTURE for K = 1 to K=5. The most likely K was K=2.

A

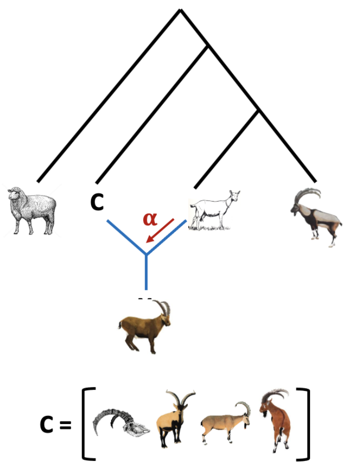


B

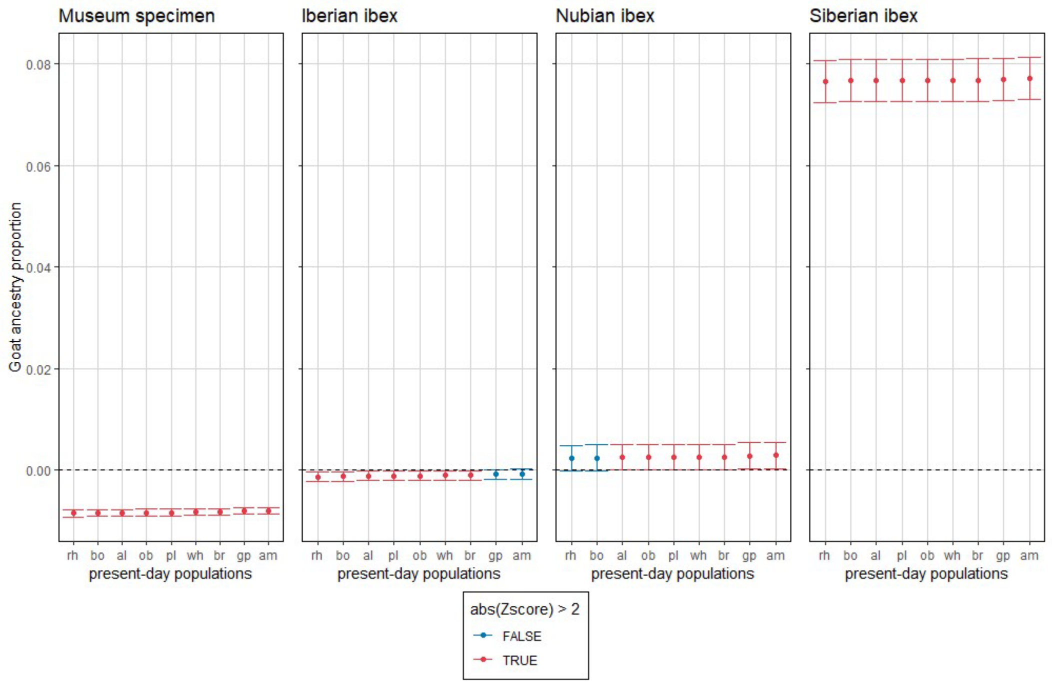


**Supplementary Figure S4: F4-ratio analysis:** A) Test design for F4-ratio analysis in admixr to estimate the proportion domestic goat ancestry in Alpine ibex depending on the sister species used (C was varied between ancient Alpine ibex, Iberian ibex, Nubian ibex and Siberian ibex). B) Proportion of domestic goat ancestry estimated as F4-ratios in admixr for 9 modern Alpine ibex populations. Each panel represents an analysis with a different outgroup. Z-score > 2 is equivalent to a p-value < 0.023 and indicates a significant divergence . Red dots and confidence intervals indicate deviance from the zero. Population abbreviations: ob - Oberbauerstock; bo - Bire Oeschinen; al - Albris; rh - Rheinwaldhorn; br - Brienzer Rothorn; wh - Weisshorn; pl - Pleureur; gp - Gran Paradiso; am - Alpi Maritimi.

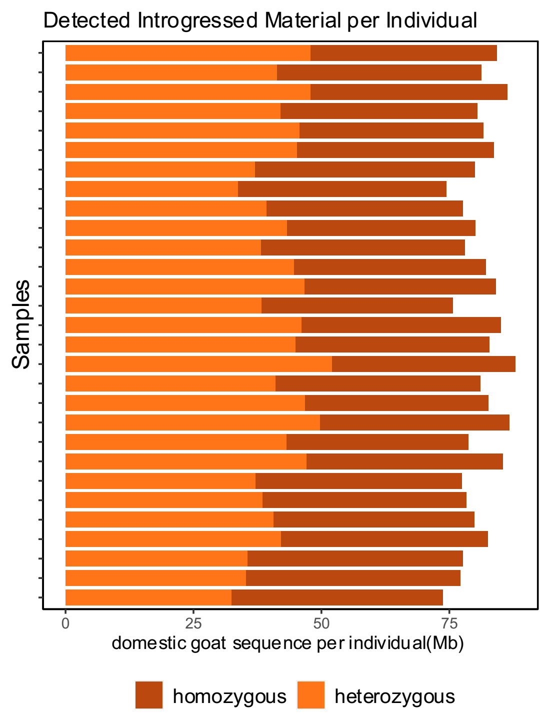


**Supplementary Figure S5:** Total introgressed sequence in Mb detected by admixfrog for each modern Alpine ibex sample. The different colors indicate homozygous (brown) and heterozygous (orange) introgressed regions, respectively.

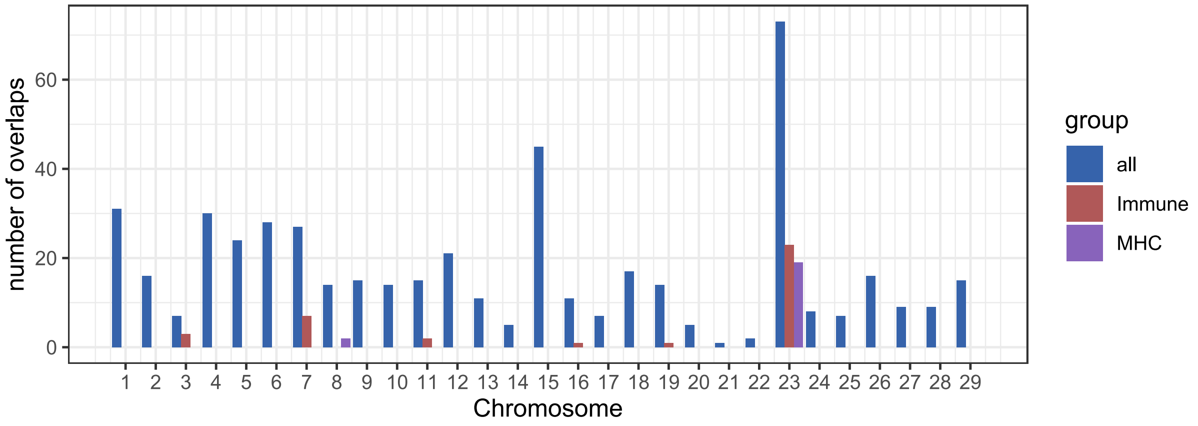


**Supplementary Figure S6:** Number of overlaps between introgressed regions identified with admixfrog and annotated coding sequences for each autosome. The different colors identify either all (red), immune associated (green) or MHC-associated (blue).

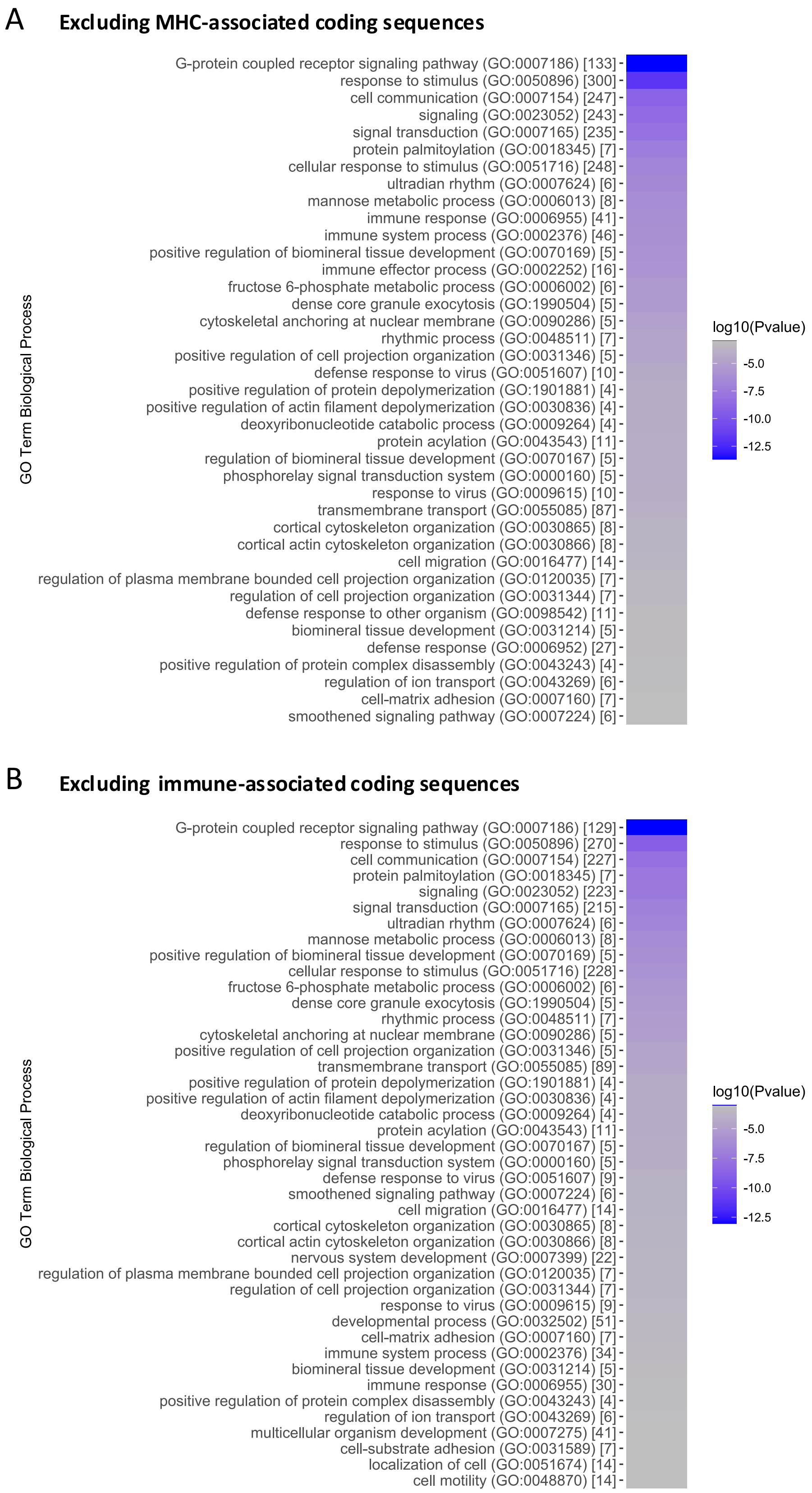


**Supplementary Figure S7:** Summary of GO term analysis for biological processes of introgressed sequences (A) excluding MHC-associated coding sequences detected by admixfrog and (B) excluding all immune-associated coding sequences detected by admixfrog. In both (A) and (B), the number in the square brackets specifies the number of genes in each category.

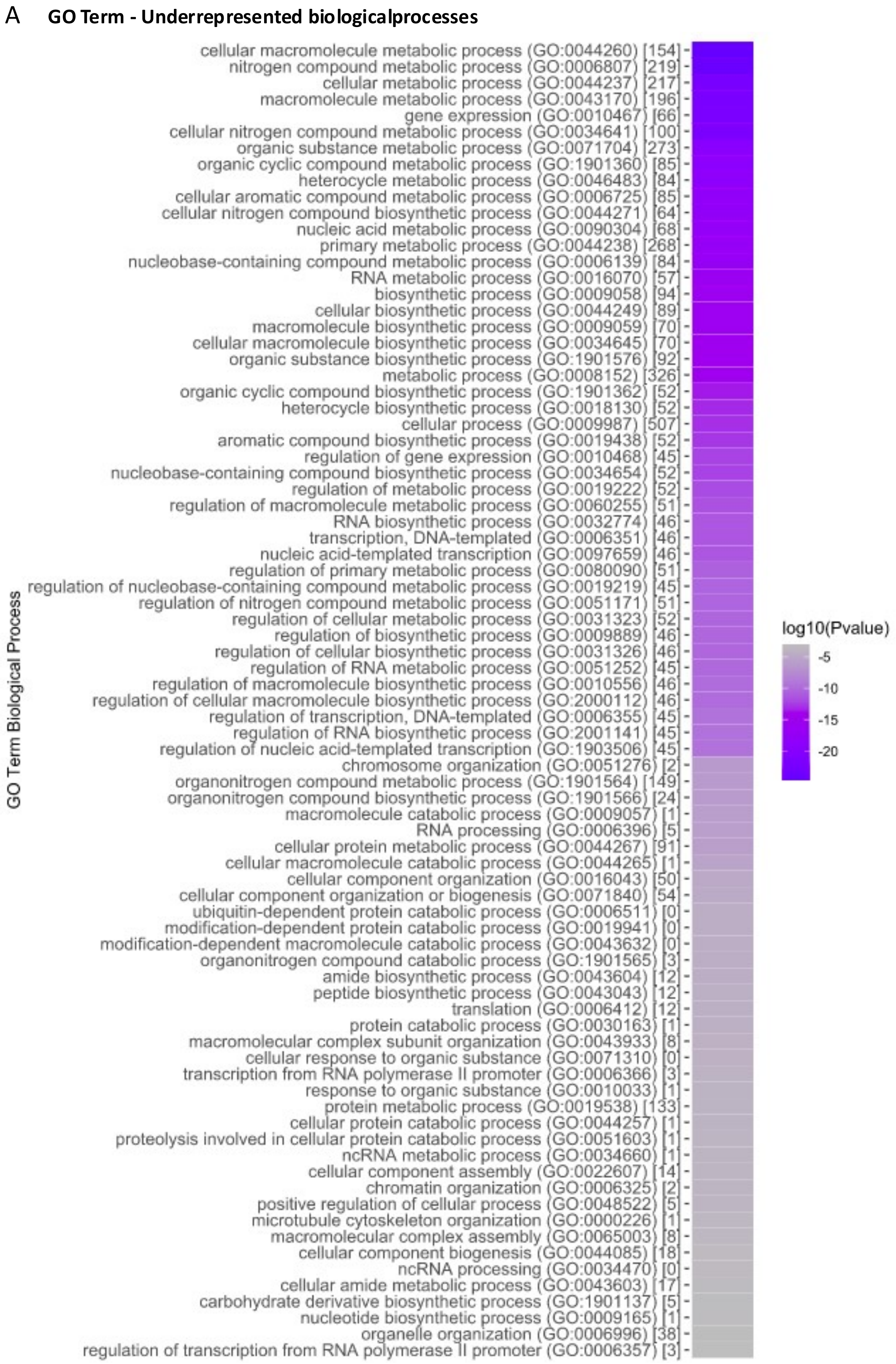


**Supplementary Figure S8:** (A) Summary of GO term analysis for biological processes with underrepresentation of introgressed sequences detected by admixfrog. The number in the square brackets specifies the number of genes in each category.

#### Supplementary Tables

**Supplementary Table 1:** Summary table describing all samples used in the analyses including location of origin and further information where available.

| Species | Sample | Population | Sex | Location |
|---|---|---|---|---|
| **Capra ibex** | ib.06S-12M_am | Alpi Maritimi | male | Rovina |
| **Capra ibex** | ib.08S-12F_am | Alpi Maritimi | female | Matto |
| **Capra ibex** | ib.22S-12F_am | Alpi Maritimi | female | Barra |
| **Capra ibex** | ib.BE0010_br | Brienzer Rothorn | female | Stelli, Hofstetten |
| **Capra ibex** | ib.BE0283_br | Brienzer Rothorn | female | Salenwang |
| **Capra ibex** | ib.BE309_bo | Bire Oeschinen | female | Schwarzhorn, Oeschinen |
| **Capra ibex** | ib.BE0407_bo | Bire Oeschinen | male | Oeschinensee, Heuberg |
| **Capra ibex** | ib.BE0463_bo | Bire Oeschinen | male | Oeschinensee, Underbärgli |
| **Capra ibex** | ib.GPO03D_gp | Gran Paradiso | male | Orco, chiapili sup |
| **Capra ibex** | ib.GPO39B_gp | Gran Paradiso | male | Orco |
| **Capra ibex** | ib.GPR21C_gp | Gran Paradiso | male | Rhemes |
| **Capra ibex** | ib.GPV05C_gp | Gran Paradiso | female | Valsaveranche, Levionaz |
| **Capra ibex** | ib.GPV09K_gp | Gran Paradiso | male | Valsaveranche, Levionaz |
| **Capra ibex** | ib.GPV13G_gp | Gran Paradiso | male | Valsaveranche, Levionaz |
| **Capra ibex** | ib.GPV18C_gp | Gran Parasdiso | male | Valsaveranche, Levionaz |
| **Capra ibex** | ib.GR0140_rh | Rheinwaldhorn | female | Scalutta Gem Safien |
| **Capra ibex** | ib.GR0201_al | Albris | female | Munt Blais, Gm S-chanf |
| **Capra ibex** | ib.GR0323_al | Albris | female | Paradies Suot, Pontresina |
| **Capra ibex** | ib.GR0442_al | Albris | male | Val da Fain Tschüffer, Pontresina |
| **Capra ibex** | ib.NW0045_ob | Oberbauerstock | male | Hoh Brisen |
| **Capra ibex** | ib.NW0061_ob | Oberbauerstock | female | Niederbauen |
| **Capra ibex** | ib.OW0002_br | Brienzer Rothorn | male | Eisee, Giswil |
| **Capra ibex** | ib.UR0036_ob | Oberbauerstock | female | Isenthal Bärenstock |

| | | | | |
|---|---|---|---|---|
| **Capra ibex** | ib.VS0031_wh | Weisshorn | female | Jungen, St. Niklaus |
| **Capra ibex** | ib.VS0079_wh | Weisshorn | female | Seetal, Grächen |
| **Capra ibex** | ib.VS0081_wh | Weisshorn | female | Jungen, St. Niklaus |
| **Capra ibex** | ib.VS0139_pl | Pleureur | male | Dixènce-Evolène |
| **Capra ibex** | ib.VS1121_pl | Pleureur | female | Verbier |
| **Capra ibex** | ib.VS1124_pl | Pleureur | male | Boussine |
| **Capra ibex** | iban.Wi1 | | | Wallis |
| **Capra ibex** | iban.Lu2 | | | Edisloch |
| **Capra pyrenaica** | py.V53_vi | | | Sierra de Gredos |
| **Capra pyrenaica** | py.V66_vi | | | Sierra de Gredos |
| **Capra pyrenaica** | py.M518_sn | Sierra Nevada | | Sierra Nevada |
| **Capra pyrenaica** | py.Z5_hb | | | Maestrazgo |
| **Capra nubiana** | nu.ibx20 | | | Sinai, Egypt |
| **Capra nubiana** | nu.ibx61 | | | Howtat, Central Saudi Arabia |
| **Capra sibirica** | si.Casi1 | | male | South of Issyk lake |
| **Capra sibirica** | si.Casi2 | | male | Dasht-i Jum Reserve, Tajikistan |
| **Capra falconeri** | fal.Cafal1 | | | Dasht-I Jum Reserve, Tajikistan |
| **Capra aegagrus aegagrus** | ae.IRCA-C3-1001 | | male | Marakan |
| **Capra aegagrus aegagrus** | ae.IRCA-G2-5063 | | male | Arasbaran |
| **Capra aegagrus aegagrus** | ae.IRCA-I11-0001 | | male | Khan Gormaz |
| **Capra aegagrus aegagrus** | ae.IRCA-I6-5237 | | male | Karnagh |
| **Capra aegagrus aegagrus** | ae.IRCA-K12-0005 | | male | Lashgar Dar |
| **Capra aegagrus aegagrus** | ae.IRCA-M12-0008 | | male | Khomein |
| **Capra aegagrus hircus** | hi.FRCH-AL-0002 | Alpine | female | |
| **Capra aegagrus hircus** | hi.FRCH-SA-0001 | Saanen | female | |

| | | | | |
|---|---|---|---|---|
| **Capra aegagrus hircus** | hi.IRCH-B3-5031 | | female | Khoiy |
| **Capra aegagrus hircus** | hi.IRCH-C5-5206 | | male | Nushin Shahr |
| **Capra aegagrus hircus** | hi.IRCH-F3-5044 | | female | Ahar |
| **Capra aegagrus hircus** | hi.IRCH-F4-5093 | | female | Vazarghan |
| **Capra aegagrus hircus** | hi.IRCH-G5-5185 | | male | Sarab |
| **Capra aegagrus hircus** | hi.ITCH-SA-0001 | Saanen | | |
| **Capra aegagrus hircus** | hi.ITCH-SA-0005 | Saanen | | |
| **Capra aegagrus hircus** | hi.MOCH-AA10-2195 | landrace | male | Maaterka |
| **Capra aegagrus hircus** | hi.MOCH-L10-3100 | landrace | male | Ayir |
| **Capra aegagrus hircus** | hi.MOCH-Q15-1143 | landrace | female | Foum Zguid |
| **Capra aegagrus hircus** | hi.MOCH-R5-0037 | landrace | female | Khmiss Sahel |
| **Capra aegagrus hircus** | hi.MOCH-S8-2252 | landrace | male | Al Hajeb |
| **Capra aegagrus hircus** | hi.MOCH-V5-3059 | landrace | female | Beni Boufrah |
| **Capra aegagrus hircus** | hi.PCGA106 | Peacock | | |
| **Ovis aries** | oa.IROA-B2-5037 | | female | Ilbolaghi, Iran |
| **Ovis aries** | oa.MOOA-AA10-2191 | Beni Guil | female | Beni Iguil, Morocco |
| **Ovis orientalis** | oa.IROO-C3-0001 | Mouflon | male | Marakan, Iran |
| **Ovis vignei** | oa.IROV-AB8-1004 | Urial | male | Dare yam, Iran |
| **Ovis canadensis** | oa.OCAN_OCAN3 | Bighorn sheep | | |

**Supplementary Table 2**: $F_3$ statistic testing if Alpine ibex represent an admixed population of the domestic goat and another group. A significantly negative value provides this evidence. However, no significant negative values are no evidence for the absence of admixture.

| A | B | C | $F_3$ | stderr | Z-Score |
|---|---|---|---|---|---|
| **Domestic goat** | Museum sample | Alpine ibex | 0.816428 | 0.011114 | 73.461 |
| **Domestic goat** | Iberian ibex | Alpine ibex | 0.919873 | 0.012703 | 72.414 |
| **Domestic goat** | Nubian ibex | Alpine ibex | 3.755556 | 0.044385 | 84.613 |
| **Domestic goat** | Siberian ibex | Alpine ibex | 4.450206 | 0.052562 | 84.665 |
| **Domestic goat** | Markhor | Alpine ibex | 10.942527 | 0.14096 | 77.629 |
| **Domestic goat** | Bezoar | Alpine ibex | 11.892547 | 0.153198 | 77.629 |
| **Domestic goat** | Sheep | Alpine ibex | 7.064663 | 0.086607 | 81.571 |
